# Dynamics of associated microbiomes during algal bloom development: to see and to be seeing

**DOI:** 10.1101/2023.09.05.556447

**Authors:** Ayagoz Meirkhanova, Adina Zhumakhanova, Polina Len, Christian Schoenbach, Eti E. Levi, Erik Jeppesen, Thomas A. Davidson, Natasha S. Barteneva

## Abstract

Our understanding of the interactions between bacteria and phytoplankton in the freshwater phycosphere, including the development of algal blooms, is very limited. To identify the taxa and compositional variation within microbial communities, we performed 16S rRNA amplicon sequencing research on samples collected weekly through summer from mesocosms that differed in temperature and mixing regimes. We investigated, for the first time, the abundance diversity of microalgae, including Chlorophyta, Cryptophyta, and Cyanobacteria species, using visualization-based FlowCAM analysis and classification of microbial communities to species level by nanopore next-generation sequencing. We found that nanopore metagenomics, in parallel with complementary imaging flow cytometry, can depict the fine temporal dynamics of microbiomes associated with visually identified *Microcystis* morphospecies, Chlorophyta, and Cryptophyta during algal bloom development. Our results showed that the temporal characteristics of microbiomes combined with a visual approach may be a key tool to predict the metacommunity structure and dynamics of algal blooms in response to anthropogenic effects and climate change.

## INTRODUCTION

Microbial interactions, including those between phytoplankton and heterotrophic bacteria, greatly impact ecosystems. Specifically, some heterotrophic bacteria, such as Alphaproteobacteria, provide autotrophic phytoplankton with essential vitamins like cobamides (the family of enzyme cofactors that include cobalamin-vitamin B12)^1–3^ in exchange for organic carbon and sulfur. They also produce siderophores that bind iron utilized by microalgae and increase the bioavailability of trace metals^7,8^, regenerate nutrients from organic materials^9,10^, and support the long-term survival of phytoplankton members^11,12^. A recent study of 332 microalgal species revealed that 54% of the analyzed algae required cobalamin, 27% required thiamin, and 8% biotin^13^. Phytoplankton-derived organic matter following algal death has a species-specific effect on bacteria^14^. The increase in the availability of different substrates induces changes in bacterial communities^15^. While significant knowledge has been accumulated on microbial-marine phytoplankton interactions^11,16–18^, comparatively less work has been published on freshwater ecosystems. Despite the importance of interactions between freshwater phytoplankton and heterotrophic bacteria, our understanding of the impacts of bacterioplankton on the phycosphere still needs to be improved^19^.

The mechanisms of how bacteria alter the phycosphere and interactions between phytoplankton and bacteria are unclear. Changes in the composition of the phytoplankton community have been shown to correlate with changes in bacterial community composition^20,21^. Algal-bacterial interactions can manifest not only as mutualistic^1,9,22,23^ but also as competition over limited inorganic resources^8,24^. Bacterial control of natural algal blooms by the *Roseobacter* lineage prevalent in algal blooms dominated by dinoflagellates and *E. huxleyi* was described in one of the early studies by Gonzalez and co-authors^25^. In freshwater systems that function as sources of drinking water for human use, it is essential to reach a detailed understanding of the control of harmful algal blooms (HAB) by bacteria^26–28^. The mesocosm experiments provided a possibility of modeling algal-microbial interactions in larger volumes and conditions closer to the natural environment than can be obtained in lab cultures ^29–32^.

Microbial communities are most commonly visualized by microscopy and flow cytometry (with cytometry processing thousands of cells per second). With few studies combining next-generation sequencing with flow cytometry and phenotyping for co-occurrence analysis of microbial communities^33^, research using imaging flow cytometry (IFC) for a search of host-microbe associations in aquatic systems is absent.

To characterize the bacterial communities associated with the freshwater phycosphere exhibiting *Microcystis*, Cryptophyta, and green algae blooms, we used integrated nanopore-based next-generation sequencing and IFC, for simplicity named mNGS-IC (microbial NGS combined with imaging cytometry). We focused on answering the following questions: (1) What are the major bacterial components of microbiomes associated with *Microcysti*s, Cryptophyta, and Chlorophyta? (2) Do the microbial communities vary with morphospecies? (3) Do bacterial communities associated with the prevalence of different *Microcystis* spp. harbor taxa that are universal, comprising a ‘core’ community? (4) How do phycosphere communities change throughout bloom development?

We hypothesize that these distinct phycosphere communities are shaped by species-specific interactions and change with variations in the physico-chemical environment (temperature and nutrients) over time. The study aimed to explore the microbial communities along the seasonal mesocosm timeline. In particular, we attempted to investigate major shifts in the bacterial community structure during *Microcystis* morphospecies, Cryptophyta, and green algal blooms modeled in a mesocosm. We suggested that initial algal bloom development would require an association with bacteria adding to the microalgae functional potential and that high abundance of algicidal bacteria would coincide in time with *Microcysti*s bloom collapse.

## RESULTS

### Microbial and phytoplankton community composition is influenced by temperature

The community composition within the mesocosms (tanks) was assessed using the mNGS-IC approach. As outlined in the experimental design, two main stages of community dynamics were achieved over the course of the experiment: stratification and mixing periods. In total, two weeks of stratification were followed by two weeks of mixing; the process was repeated twice. Weeks 1, 2, 5, and 6 corresponded to stratification periods, and weeks 3, 4, 7, and 8 to mixing. Samples were taken at two depths – surface and bottom – during the whole duration of the experiment. Changes in temperature and oxygen levels are outlined in **Suppl. Figure 1**. An overall decrease in temperature was observed in all tanks throughout the eight weeks of the experiment, reflecting declining temperatures in Denmark over the summer period (**Suppl. Figure 1**). Notable differences in the temperature of the surface and bottom layers were recorded during the first stratification period; similarly, oxygen levels were generally higher in the surface water than in the bottom layers during both stratification periods, indicating that varying levels of stratification were achieved in the tanks.

The temperature in tank D1, with an ambient temperature regime, ranged from 16°C to 25.2°C and was similar in the surface and bottom layers throughout the experiment. The exceptions were weeks 1 and 2 (corresponding to the first stratification period), where temperatures in the bottom layer were slightly lower than in the surface layer. More notable stratification between the layers was observed for the oxygen levels, which ranged from 11.41 to 19.37 mg/L in the surface layers and from 0.07 to 14.88 mg/L in the bottom layers. The mixing periods were characterized by relatively similar oxygen levels between the layers, while oxygen during the stratification periods was generally found to be lower in the bottom layers than in the surface. Both sequencing and imaging cytometry-based analyses revealed dominance of cyanobacteria (ranging from 27.2% up to 68.8% based on 16S and from 85.7% up to 99.2% based on IFC) throughout the experiment in the phycosphere of tank D1 (**Figure 1**). The 16S-based analysis identified 52 cyanobacterial species in the tank, with *M. aeruginosa* being the dominant one. In parallel, using the IFC-based approach, it was possible to identify five *Microcystis* morphospecies: *M. novacekii, M. ichtyoblabe, M. smithii, M. aeruginosa,* and *M. wesenbergii*. Following cyanobacteria, proteobacteria comprised the second largest bacterial group in this tank (14.9% up to 63.5%), with 618 species identified. Apart from cyanobacterial and proteobacterial phyla, Bacteroidetes comprised the third largest phylum in this tank, with a relative abundance ranging from 5.4% to 31.9% and 222 species identified.

**Figure 1.**
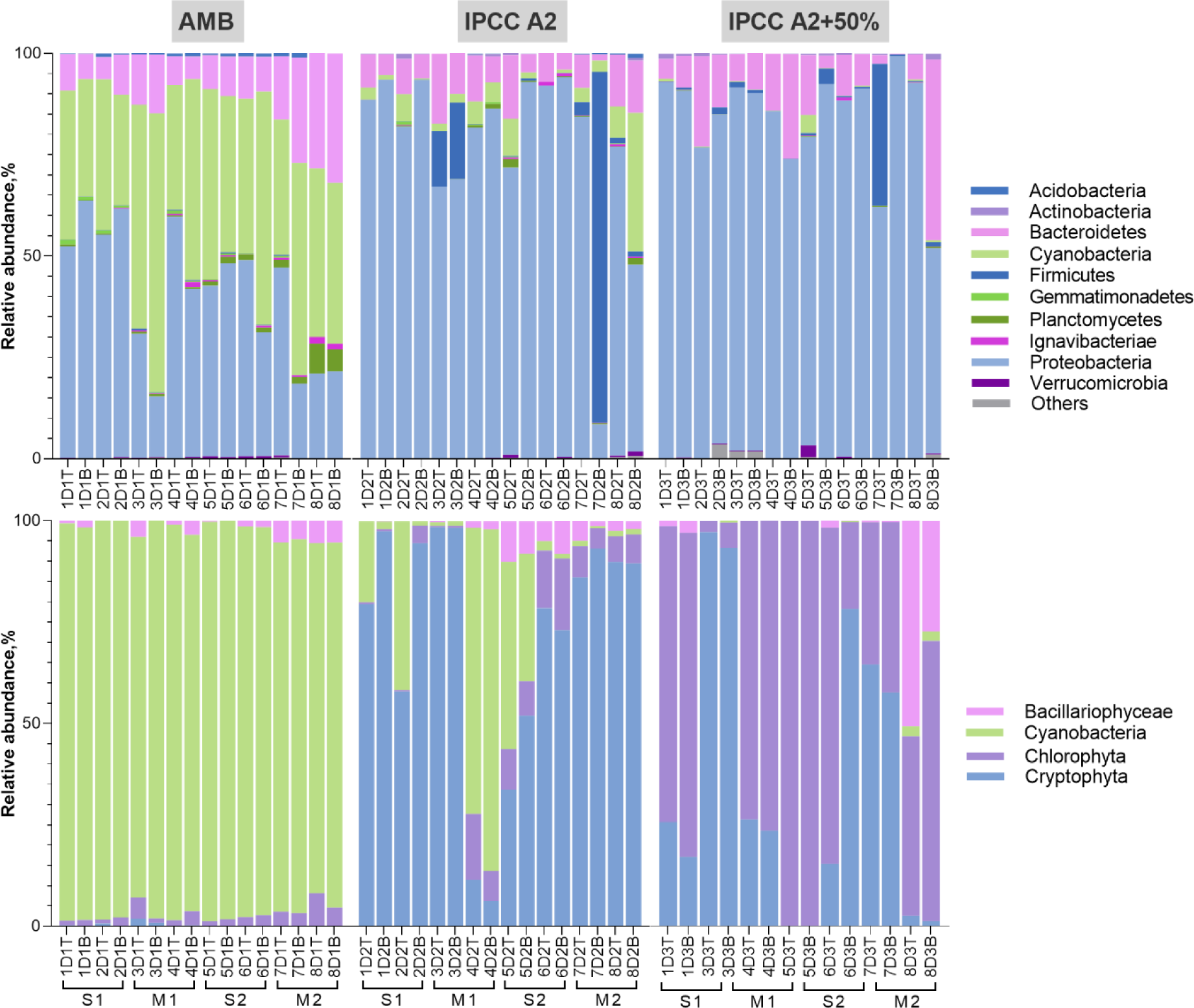
Relative abundances of dominant bacterial phyla (**top row**) and phytoplankton groups (**bottom row**) across three temperature regimes (AMB, IPCC A2, IPCC A2+50%). S1 and S2 – first (weeks 1 and 2) and second (weeks 5 and 6) stratification periods respectively, and M1 and M2 – first (weeks 3 and 4) and second (weeks 7 and 8) mixing periods respectively. Labels “T” and “B” on the x-axis represent surface and bottom samples respectively.

Tank D2, had an elevated temperature regime (IPCC A2), ranging from 18.8°C to 33.1°C. Similar to tank D1, temperature-based stratification was observed during the first stratification period (weeks 1 and 2), and notable differences in oxygen levels between the surface and bottom layers were recorded during both stratification periods. 16S-based analysis revealed dominance of the proteobacteria phylum, whose relative abundance ranged from 8% to 93.8% (**Figure 1**) during the experiment. Bacteroidetes were the second largest phylum identified, with abundances varying from 1.4% to 17.2%. In addition, both 16S- and IFC-based analyses revealed presence of cyanobacteria in tank D2. According to NGS results, the peak relative abundance of the cyanobacterial phylum was observed during the end of the second mixing period (week 8) - 34.2%, with Nostocaceae being the dominant family. IFC-based analysis revealed slightly different temporal dynamics of cyanobacteria abundance, with a maximum peak during week 4 corresponding to a slight temperature increase prior to the sampling date (**Suppl. Figure 1**). Unlike tank D1, in which the cyanobacteria group was predominantly composed of several *Microcystis* morphospecies, *Aphanocapsa* sp. was found to be the major contributor to the cyanobacterial abundance in tank D2. Lastly, the analysis of major phytoplankton groups revealed a large proportion of the Cryptophyta phylum with varying dominance throughout the experiment. *Cryptomonas* sp. was identified as the dominant genus within this group. Peak Cryptophyta abundance was found during the beginning of the first mixing period (week 3) in both layers- 98.8% (6,793 particles/ml) on surface and 98.9% (6,839 particles/ml) in the bottom layer, while its minimum (6%) corresponded to the increase of cyanobacteria during week 4.

Lastly, notable differences in temperature and oxygen levels between surface and bottom layers of tank D3 (IPCC A2+50% temperature regime) during stratification periods were recorded. The temperature in the surface layers ranged from 20.3°C to 31.5°C and from 20.1 °C to 23.9 °C in the bottom, while oxygen levels ranged from 0.1 to 26.6 mg/L (surface) and from 0.03 to 12.7 mg/L (bottom). Similar to tank D2, proteobacteria comprised the largest bacterial phylum in tank D3, with relative abundances ranging from 62.2% to 92.9% in the surface and from 50.7% to 99.4% (**Figure 1**) in the bottom layers. Parallel IFC-based analysis found Chlorophyta to be the dominant phytoplankton group in this tank (with the exception of weeks 2 and 7), and the group was mainly composed of *Micractinium* sp. The beginning of the second stratification period (week 5) corresponded to peak Chlorophyta abundance - 99.9%. In addition to Chlorophyta, Cryptophyta was the second largest phylum identified using the IFC-based approach, with maximum relative abundance of 97.1% during week 3. Cyanobacteria presence was minimal in this tank.

Apparent clustering of samples based on temperature regime was observed using non-metric multidimensional scaling (NMDS) analysis for both 16S- and FlowCAM-based analyses (**Figure 2**). Distinctive bacterial and phytoplankton communities were formed at varying temperatures (AMB, IPCC A2, and IPCC A2+50%) over the course of the experiment. Pairwise analysis of similarities (ANOSIM) revealed a significant difference between these communities in tanks D with different temperature regimes (p-value<0.05).

**Figure 2.**
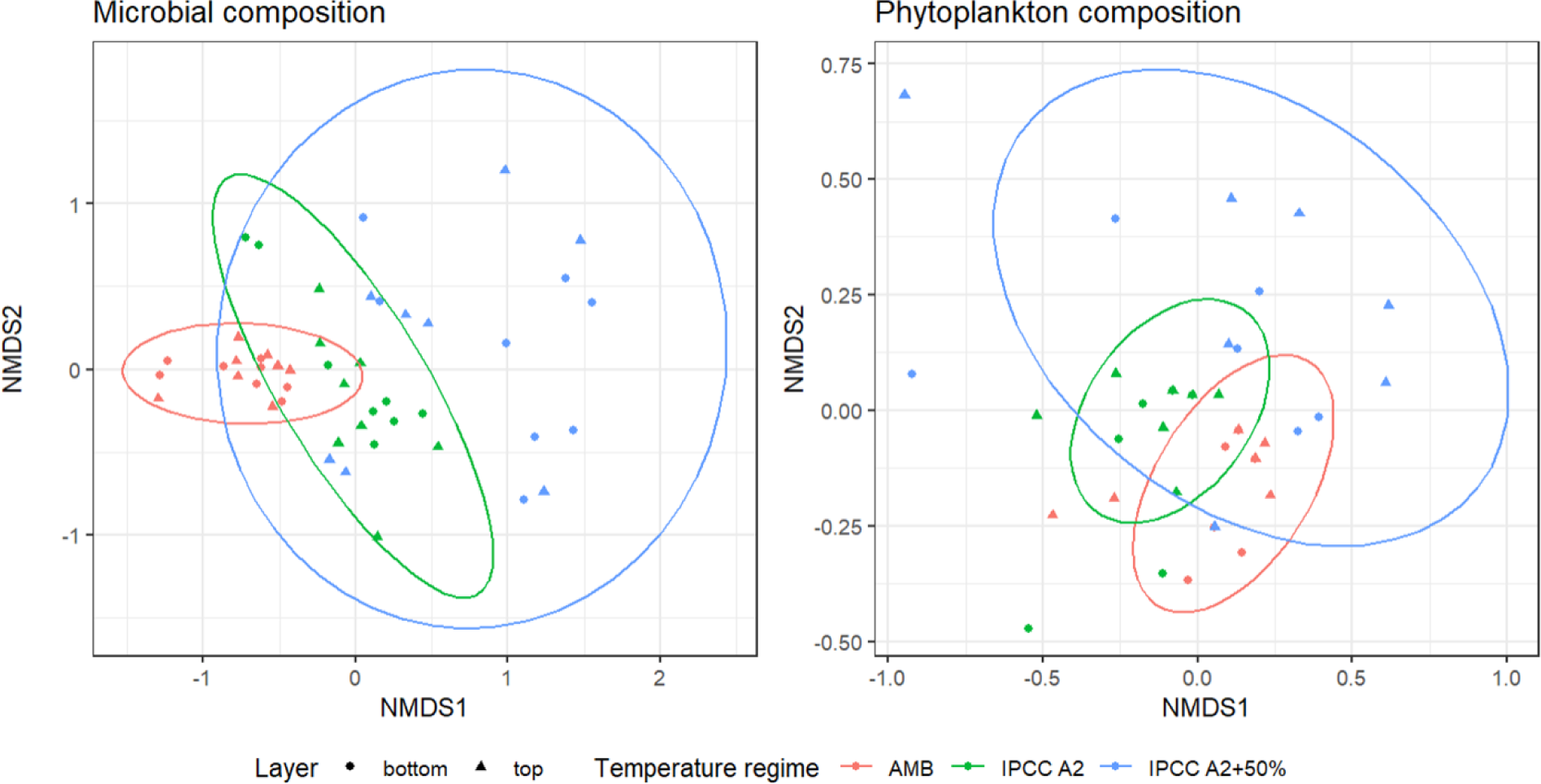
Non-metric multidimensional scaling (NMDS) analysis of community compositions throughout the experiment. NMDS ordination plots with stress values of 0.16 (left) and 0.14 (right) indicate clustering of microbial and phytoplankton communities across temperature regimes. Each ellipse and color represent a different treatment group: AMB – red, IPCC A2 – green, and IPCC A2+50% – blue. Ranked dissimilarities between all identified clusters were significantly different from each other for both microbial and phytoplankton compositions (R=0.54, p-value=0.001 and R=0.34, p-value=0.001, respectively).

### *Microcystis*-associated microbial clusters

As mentioned, five *Microcystis* morphospecies were identified using FlowCAM-based cytometric analysis (*M. novacekii, M. ichtyoblabe, M. smithii, M. aeruginosa, M. wesenbergii*)^34^. However, morphospecies that were difficult to identify, namely *M. smithii* and *M. aeruginosa*, were grouped into a single cluster. The identified *Microcystis* species were then divided into two main groups: colonial (consisting of five morphospecies) and non-colonial small clusters (NCSC). **Figure 3 (B, C)** illustrates changes in the absolute abundance of these two groups in the surface and bottom layers of tank D1. A peak of NCSC was observed in the surface layers (60,847 particles/ml) during the beginning of the second stratification period (week 5). In addition, an overall drop of NCSC *Microcystis* during the beginning of the second mixing period (week 7) in the surface layer (5,110 particles/ml) was accompanied by a decrease in its abundance in the bottom layer as well (from 14,959 particles/ml during week 7 to 5,685 particles/ml by the end of the experiment). Overall, the abundance of non-colonial *Microcystis* had decreased by the end of the experiment, with an increasing number of colonial forms.

**Figure 3.**
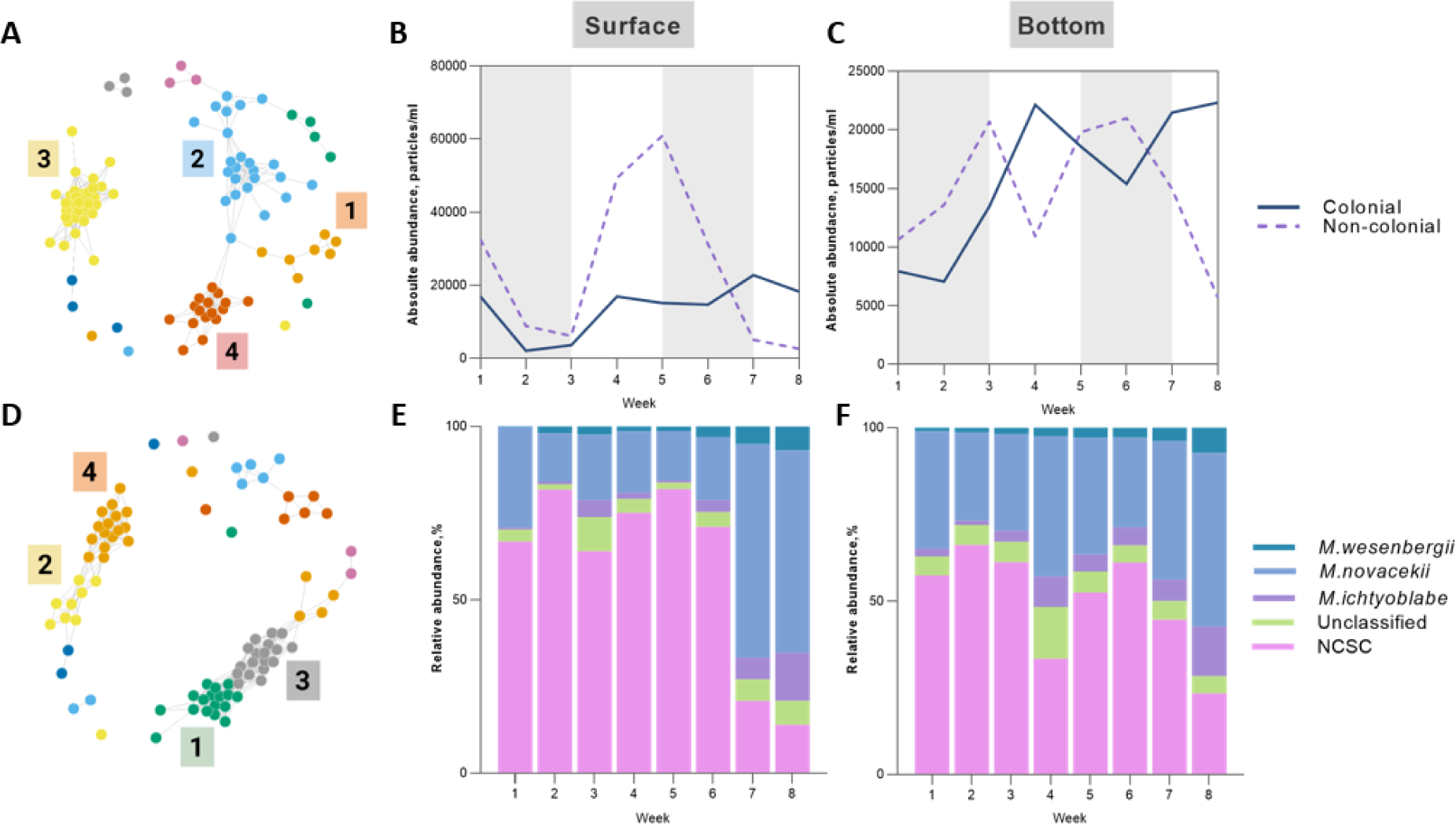
*Microcystis-*associated microbial clusters. (**A, D**) Co-occurrence network based on Pearson correlation coefficient (R>0.75, p-value<0.05) for surface samples – (**A**) and bottom samples (**D**) of tank D1, four major microbial clusters were identified in each layer and color-coded separately for each cluster; (**B, C**) absolute abundance of colonial and non-colonial (NCSC) forms of *Microcystis* throughout the experiment in surface and bottom layers, shaded areas correspond to stratification periods; (**E, F**) relative abundances of *Microcystis* morphospecies and non-colonial forms throughout the experiment in surface (**E**) and bottom (**F**) layers.

The dynamics of separate morphospecies are depicted in **Figure 3 (E, F)**, and varying dominance of *M. novacekii,* the most abundant morphospecies during the experiment, was observed in both layers. Maximum peak of *M. novacekii* abundance was recorded during week 7 – at the beginning of the second mixing period – in the surface layer (15,209 particles/ml) and in the bottom layer (13,411 particles/ml). *M. wesenbergii* and *M. ichtyoblabe* contributed almost equally to the total abundance of *Microcystis*.

Correlation analysis based on the Pearson correlation coefficient was conducted to determine microbial species co-occurring throughout the experiment. Reads were first filtered to eliminate species with <1% relative abundance; obtained correlation coefficients less than 0.75 were filtered out (p-value < 0.05) prior to network construction. Resultant networks in tank D1 (**Figure 3A, D**) revealed eight microbial clusters with the largest number of members – four clusters for the surface and four for the bottom layers. The composition of each cluster is listed in **Tables 1** and **2**.

The Pearson correlation coefficient-based matrix revealed significant (p-value<0.05) correlations between some of the microbial clusters identified in the network analysis and *Microcystis* morphospecies. A significant positive correlation was identified between microbial cluster 3 and *M. ichtyoblabe* abundance (Pearson coefficient r= 0.81). Peak *M. ichtyoblabe* abundance was recorded during week 8, corresponding to the peak in the abundance of cluster 3 (**Figure 4A**). Cluster 3 consisted of different members of the Bacteroidetes, cyanobacteria, and proteobacteria phyla (**Table 1**). Sphingobacteriales were the largest order within this cluster, with 12 species identified. In addition to this, a significant negative correlation (r= -0.71) was found between cluster 4 and the cumulative abundances of *M. smithii* and *M. aeruginosa* morphospecies in the surface layers of tank D1 (**Figure 4B**). Cluster 4 mainly encompassed members of the Alphaproteobacteria class, namely the Rhodospirillales order (**Table 1**). The main peak of this cluster corresponded to the decrease in the absolute abundance of *M. smithii* and *M. aeruginosa* morphospecies during the end of the first stratification period (week 2). Vice versa, the maximum abundance of *Microcystis* morphospecies during week 4 was accompanied by a decline in the abundance of the microbial cluster 4.

**Figure 4.**
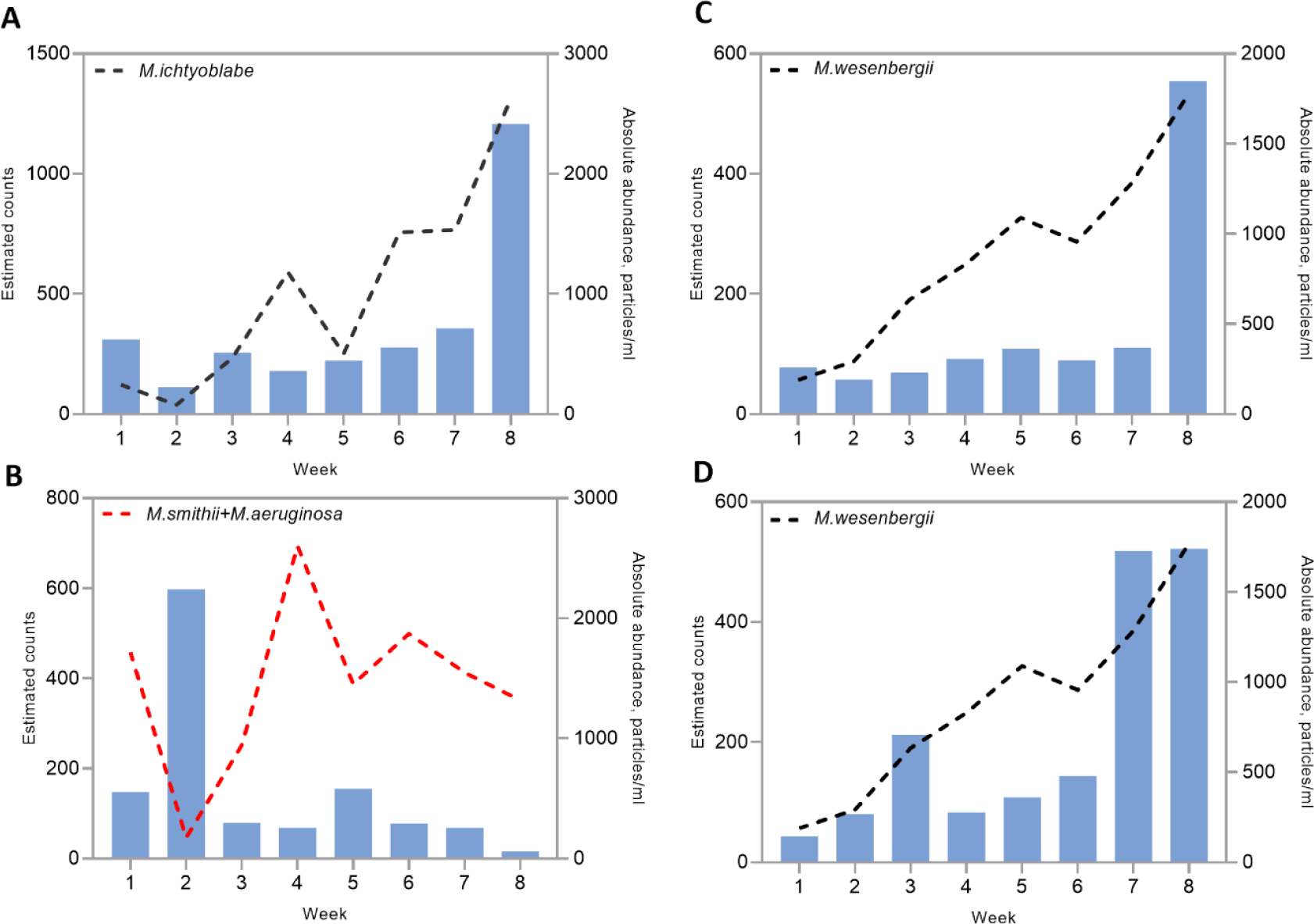
Dynamics of *Microcystis* spp. abundance against significantly (p-value<0.05) correlated microbial clusters in tank D1. Blue bars represent abundance of each significantly correlated microbial cluster. (**A**) *M. ichtyoblabe* absolute abundance against positively correlating microbial cluster 3 in the surface layers; (**B**) *M. smithii* and *M. aeruginosa* group absolute abundances against negatively correlating microbial cluster 4; (**C**) *M. wesenbergii* absolute abundance against positively correlating microbial cluster 1 in the bottom layers; (**D**) *M. wesenbergii* absolute abundance against positively correlating microbial cluster 3 in the bottom layers. Negative correlation is shown with red dashed line. Mind the different y-axis scales.

**Table 1.**
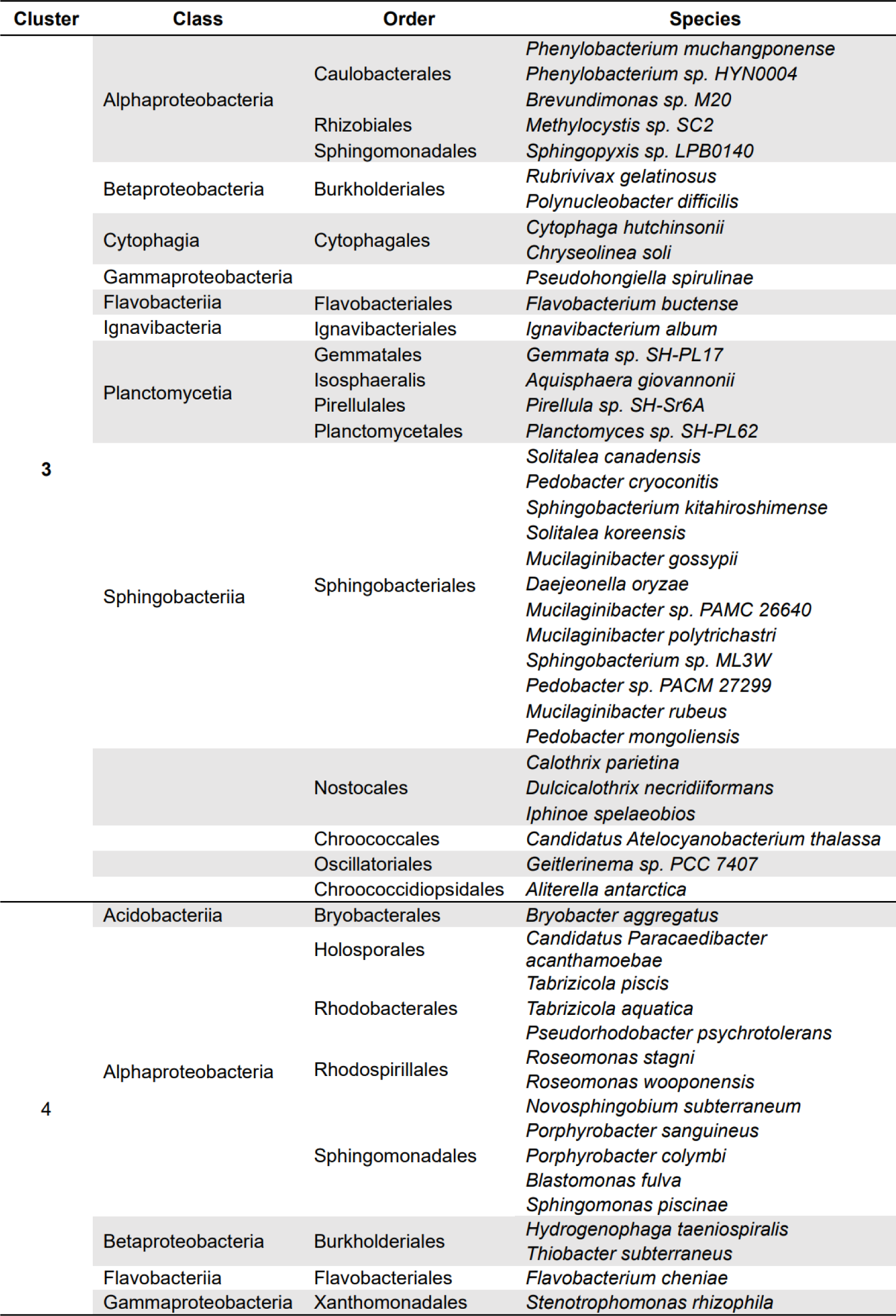
The species composition of microbial clusters with significant correlations (p-value<0.05, r>|0.7|) with *Microcystis* morphospecies in the surface layers of tank D1.

Positive correlations (p-value<0.05) between the abundance of *M. wesenbergii* and identified microbial cluster were also found in the bottom layers of tank D1. In particular, positive correlations with microbial clusters 1 and 3 were identified (r= 0.76 and 0.81, respectively) (**Figure 4C, D**). Interestingly, the members of clusters 1 and 3 of the bottom layer (**Table 2**) were similar to the members of cluster 3 in the surface layer, which had a significant positive correlation with *M. ichtyoblabe* abundance. The peaks of clusters 1 and 3 occurred together with a peak in the abundance of *M. wesenbergii* during week 8. Cluster 1 was composed of members of proteobacteria, Planctomycetes, cyanobacteria, and Bacteroidetes phyla, while the composition of cluster 3 was dominated by Bacteroidetes, with minor contributions from proteobacteria, Planctomycetes, and cyanobacteria. A difference between the clusters was observed during weeks 3 and 7 (beginning of mixing periods), where the abundance of cluster 3 increased.

**Table 2.**
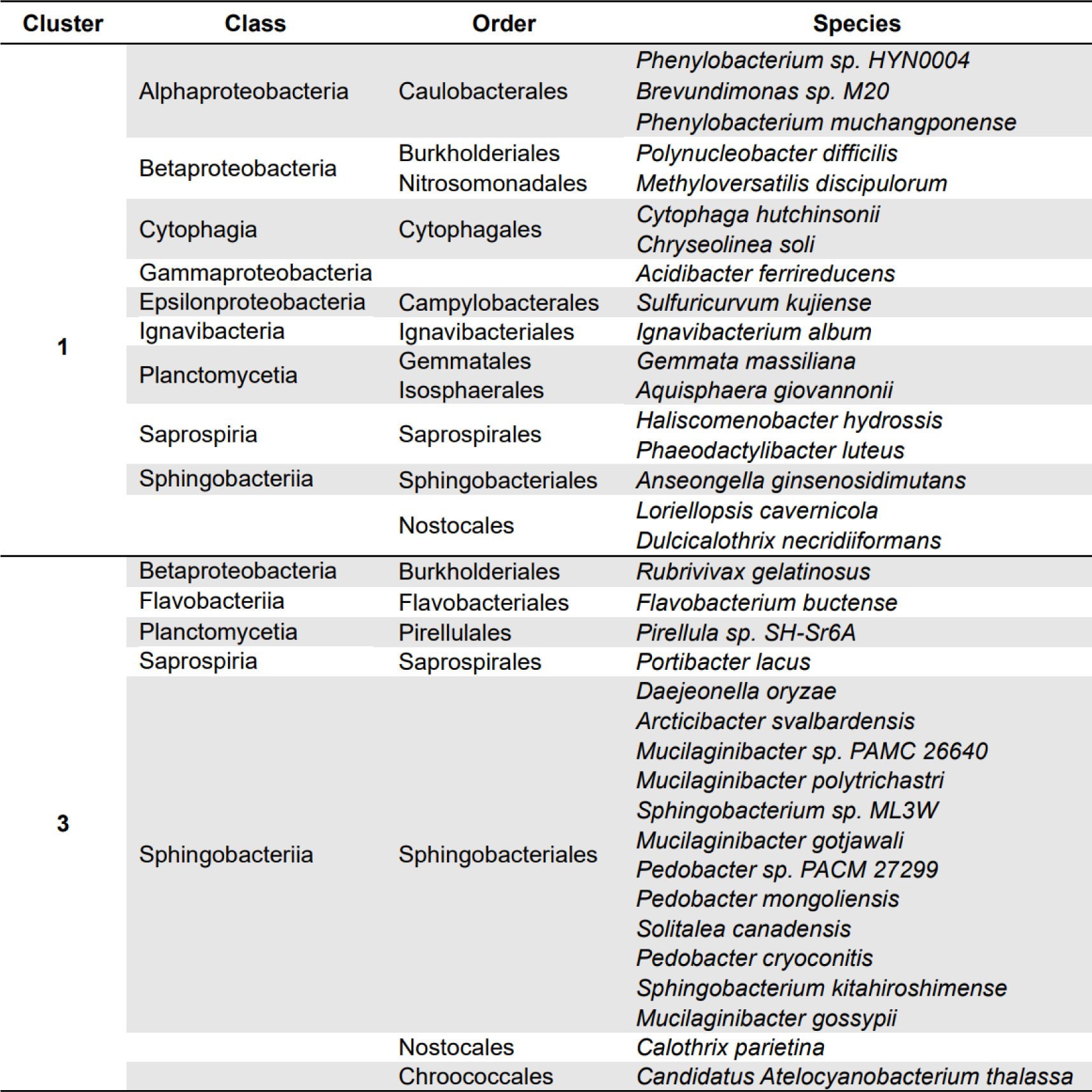
The species composition of microbial clusters with significant correlations (p-value<0.05, r>0.75) with *Microcystis* morphospecies in the bottom layers of tank D1.

### Cryptophyta-associated microbial clusters

Cryptophyta phylum members mainly dominated tank D2 with an IPCC A2 temperature regime. Dominance varied throughout the experiment (**Figure 5**); a maximum peak of Cryptophyta abundance in the surface layer occurred at the beginning of the first mixing period (98.8% in the surface layer and 98.2% in the bottom layer). Correlation analysis based on the Pearson correlation coefficient was conducted to construct networks for bacterial clusters with similar co-occurrence patterns (**Suppl. Figure 2A,B**). The resultant networks were correlated with the absolute abundance of identified Cryptophyta representatives. The data obtained indicated a significant positive correlation in the surface layer between the Cryptophyta phylum and *Massilia aurea* (r = 0.82, p-value<0.05), a member of the Betaproteobacteria class. Common peaks between the two groups were found during weeks 1, 3, and 7. In addition, a microbial cluster negatively correlating (r = -0.78, p-value<0.05) with the Cryptophyta phylum was identified in the surface layer of tank D2. This cluster comprised members of the Betaproteobacteria, Alphaproteobacteria, and Chitinophagia classes, *Limnohabitans* sp.63ED37-2 being the most abundant species. The highest abundance peaks of this cluster occurred together with sharp declines in the abundance of Cryptophyta and were observed during the end of the first stratification period (week 2) and the second mixing period (weeks 4-6).

**Figure 5.**
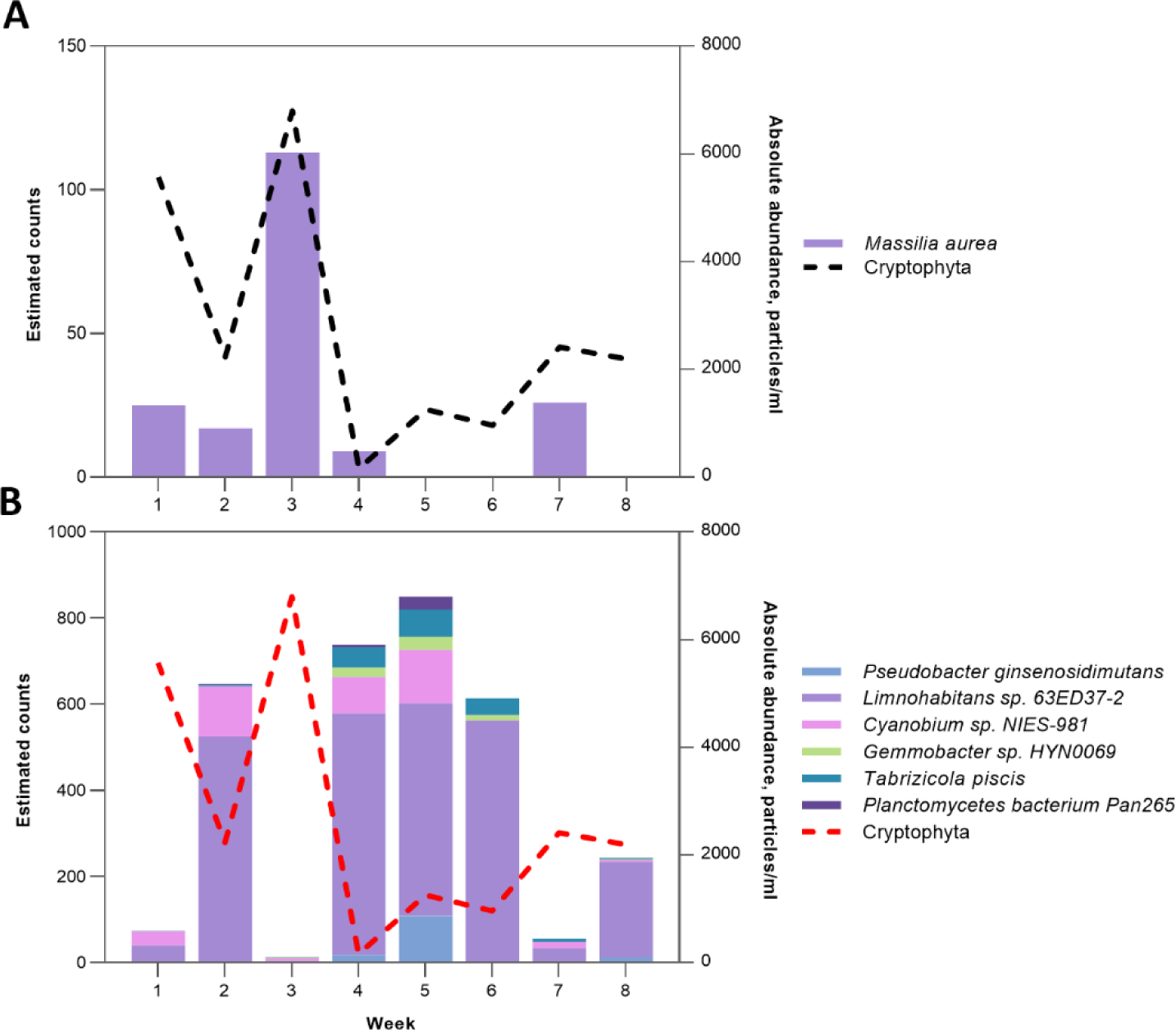
Dynamics of the Cryptophyta phylum and associated bacterial species. (**A**) Cryptophyta absolute abundance against positively correlating (p-value<0.05) *Massilia aurea* in the surface layers of tank D2; (**B**) Cryptophyta absolute abundance against negatively correlating (p-value<0.05) microbial cluster in the surface layers of tank D2. Negative correlation is shown with red dashed line. Mind the different y-axis scales.

### Chlorophyta-associated microbial clusters

Tank D3, with the IPCC A2+50% temperature regime, was dominated by members of the Chlorophyta phylum. Using FlowCAM-based IFC, three main representatives were identified: *Pediastrum* sp., *Scenedesmus* sp., and *Micractinium* sp., with *Micractinium* sp. being the most abundant genus. The relative abundance of Chlorophyta in tank D3 varied from 2.7% to 99.9% in the surface waters and from 6.2% to 99.9% in the bottom layer, with a peak reached during week 5 (the beginning of the second stratification period) (**Figure 6**). No significant differences were observed in the abundance between the surface and bottom layers. Similar to the previously outlined approach, the Pearson correlation coefficient was calculated to construct microbial co-occurrence networks and further identify any significantly correlating microbial clusters against the Chlorophyta phylum (**Supplementary** Figure 2C,D). **Figure 6** demonstrates the identified correlations. Significant positive correlations (r = 0.96) were discovered between Chlorophyta and one of the microbial clusters in the surface layers of tank D3. The maximum peak in the abundance of the microbial cluster corresponds to the peak in the abundance of the Chlorophyta phylum. The cluster was composed of members of the Betaproteobacteria class, namely *Ferribacterium limneticum, Dechloromonas aromatica*, and *Zoogloea caeni,* along with microbes of the Verrucomicrobiae and Chitinophagia classes. In addition, significant positive correlations were identified in the bottom layers of tank D3 between Chlorophyta and one of the microbial clusters (r = 0.95). Similarly, the peaks in the abundance of both groups occurred in week 5 of the experiment. This cluster was composed of four members of the Gammaproteobacteria class (*Pseudomonas syringae, Stenotrophomonas maltophilia, Stenotrophomonas rhizophila*, and *Stenotrophomonas sp. MYb57*), the Betaproteobacteria class (*Pseudoduganella danionis* and *Massilia armeniaca*), and the Bacilli class (*Exiguobacterium sp. U13-1*).

**Figure 6.**
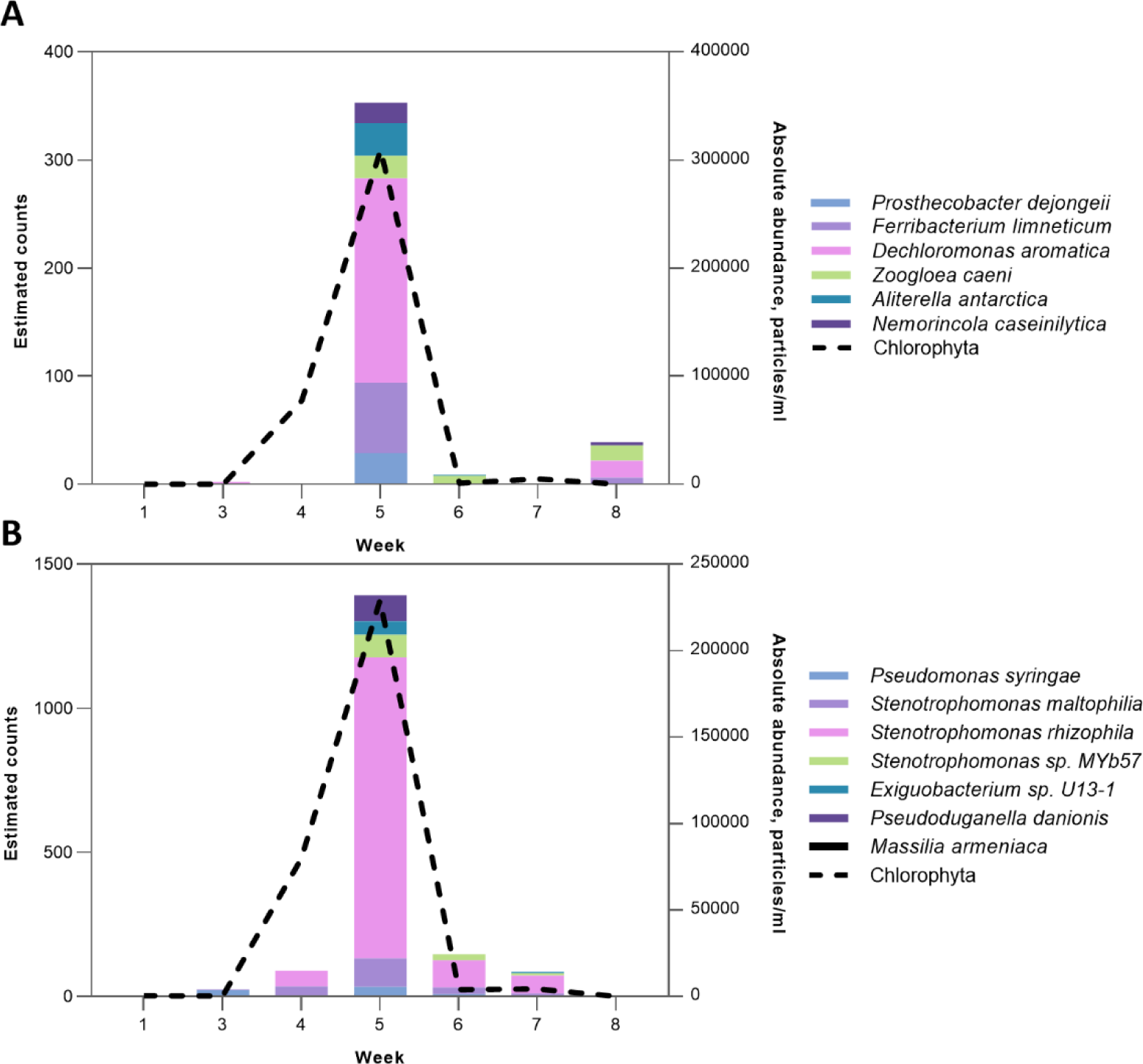
The dynamics of the Chlorophyta phylum and associated bacterial species. (**A**) Chlorophyta absolute abundance against the positively correlating (p-value<0.05) microbial cluster in the surface layers of tank D3; (**B**) Chlorophyta absolute abundance against the positively correlating (p-value<0.05) microbial cluster in the bottom layers of tank D3. Mind the different y-axis scales.

### Temporal changes in alpha diversity of microbial communities

To evaluate the changes in biodiversity within the mesocosm tanks across eight time points, variations in the Shannon index and the observed species richness for the microbial communities were calculated and presented in **Figure 7**. An overall drop in species richness was observed for both the surface and bottom layers in tank D1 (**Figure 7A**). A decrease in the Shannon index was observed during week 3, simultaneously with cyanobacteria dominance. A similar pattern was observed for weeks 6 and 7 in the bottom layers. Richness in both layers of tank D2 had an opposite trend, with an overall increase in species richness (**Figure 7B**). The increase in the proportion of the Firmicutes phylum was reflected in the decreased Shannon index during week 7 for the bottom layer. **Figure 7C** illustrates changes in the diversity indices in tank D3, where an increase in both indices during week 5 in the surface water layer occurred simultaneously with a rise in the total number of Chlorophyta representatives.

**Figure 7.**
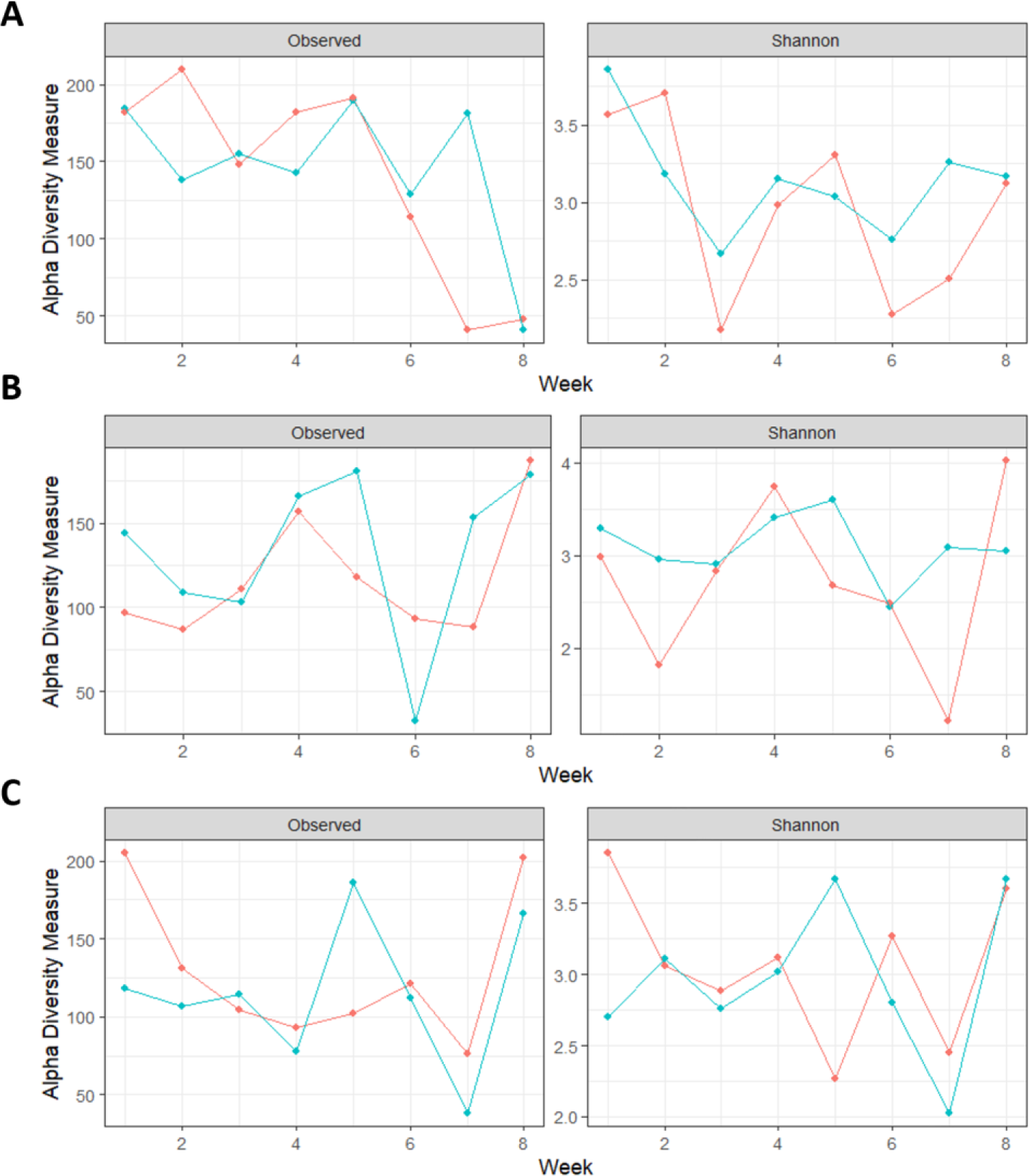
Temporal variability of microbial communities throughout the experiment for the observed species richness and Shannon index in tank D1 – (**A**), D2 – (**B**), and D3 – (**C**). Blue lines indicate surface samples and red bottom samples.

### CCA analysis of environmental gradients

Canonical correspondence analysis (CCA) was conducted to analyze the environmental gradients and their effect on community composition. The factors included in the analysis were pH, oxygen levels, temperature, stratification index, total phosphorous (TP), and PO_4_P levels. The obtained CCA plot is shown in **Figure 8**. The overall model, as well as the first two CCA axes, were statistically significant. However, constrained axes explained only 1.50 out of 6.62 of the total inertia (22.7%).

**Figure 8.**
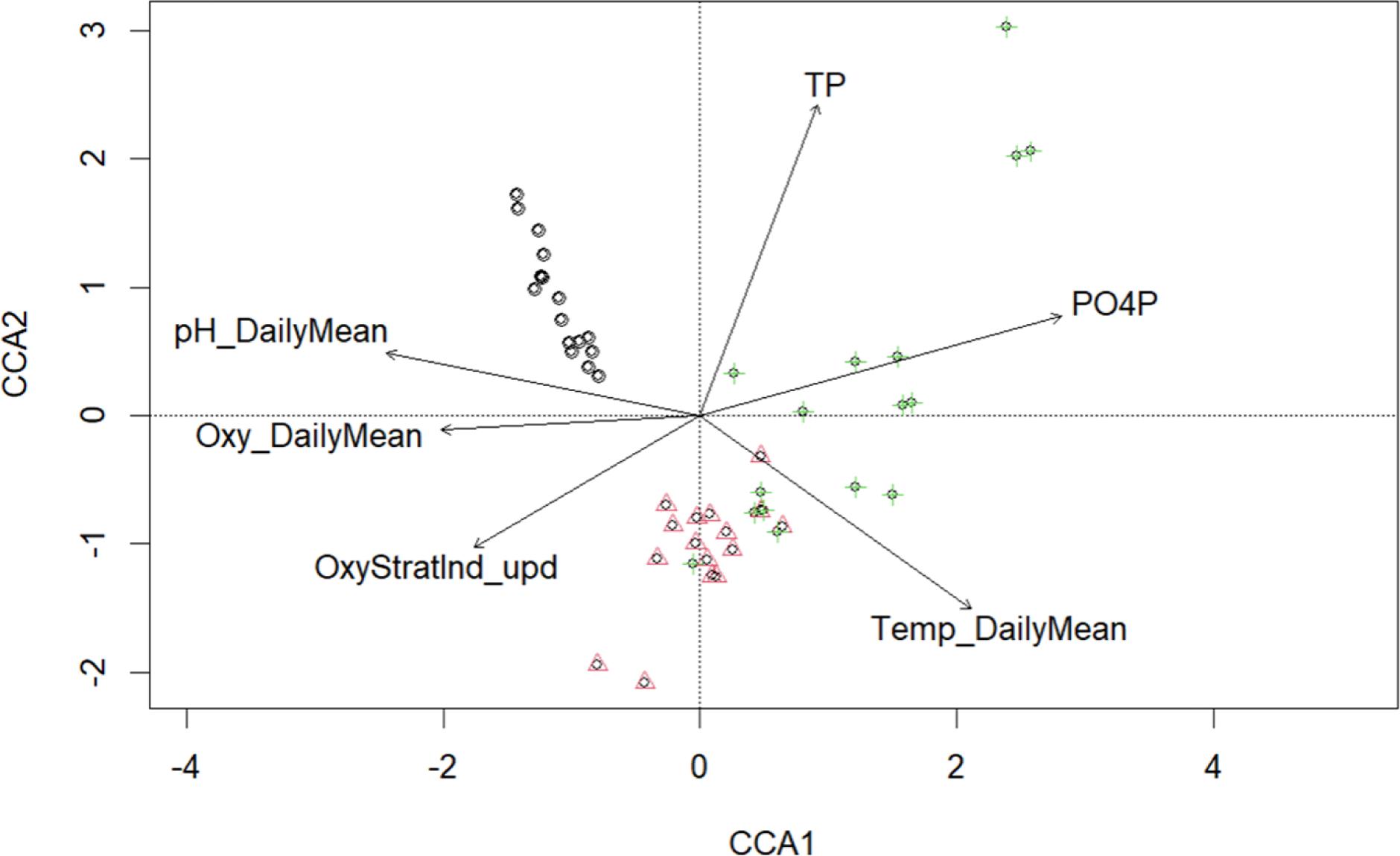
Canonical correspondence analysis (CCA) of microbial community composition and selected environmental parameters. Black color corresponds to samples from different dates with an ambient temperature regime, red to the IPCC A2 regime and green to the IPCC A2+50% regime. (pH_DailyMean – pH levels, Oxy_DailyMean – oxygen levels, OxyStratInd_upd – stratification index, Temp_DailyMean – temperature, TP – total phosphorous, PO4P - phosphate).

## DISCUSSION

The microalgae-microbiome interactions are central in natural aquatic ecosystems and in artificial cultures. However, few studies have focused on elaborating a detailed characterization of algal-associated microbes^35^; most existing studies address the interactions of microbiomes with microalgae at class and genera levels^36^. The spatiotemporal dynamics of bacterial communities and individual bacterial ecotypes are essential to understanding bacterial-algal community composition and interactions. There are “generalist” bacterial species, which occur throughout the whole season and “specialists”, which appear in significant numbers only for a limited amount of time or irregularly^37^.Heterotrophic bacteria in freshwater ecosystems are responsible for most organic matter cycling and for a significant part of the system respiration^38^. The co-occurring heterotrophic bacteria could have either negative or positive impacts on algal blooms, contributing to nutrient cycling, phytoplankton, and the lysis and degradation of toxins^39^. Field and laboratory studies revealed bacterial clusters attached to cyanobacterial and microalgal aggregates, creating an algal microenvironment or phycosphere^40^and synchronization between planktonic bacteria growth with phytoplankton bloom^41,42^.

Sequence analysis of rRNA genes ^43,44^ and comparative analysis of marker genes^45^ have revealed a distinct set of “freshwater-specific” bacterial taxa with consistent temporal differences in the composition complexity of bacterial communities’^46^. Furthermore, most microbial community studies focus on either bacterial or eukaryotic communities, but investigations aimed at obtaining an integrated view of the temporal dynamics of changes, eventually revealing the underlying ecological inferences, are scarce^47,48^, in particular for freshwater ecosystems^49,50^. An emerging approach to microbial community research based on NGS is “correlation networks”, which can be used to determine drivers in environmental ecology and help researchers in hypothesis generation ^47,51,52^. Recently, single *Microcystis* colonies microbiomes and *Microcystis*-epibiont communities were analyzed^53–55^. It was found that *Microcystis* blooms are accompanied by a diverse community of heterotrophic bacteria that play an important role in cyanobacterial bloom development and duration^54,55^. However, the molecular techniques used to characterize the *Microcystis*-associated microbiomes need more resolution in identifying the member bacteria at species level^53^. In the present study, we applied nanopore-based NGS analysis of 16S amplicons and visualization-based IFC to simultaneously characterize the dynamics of diversity and co-occurrence in two domains of life, bacteria, and Eukarya, in mesocosms with different regimes of temperature and mixing.

### *Microcystis*-associated microbiomes

*Microcystis* is characterized by great phenotypic plasticity, and currently > 50 *Microcystis* species have been identified by microscopy, often being referred to as morphospecies, including *M. aeruginosa, M. flos-aquae, M. ichthyoblabe, M. wesenbergii, M. novacekii* and others^56,57^. However, a comparison of 16S rRNA species has revealed >99% similarity and inconsistency of physiological and genetic analysis, suggesting that some morphospecies represent a single species ^34,58^. Lately, whole genome sequencing and phylogenetic clustering intended to differentiate *Microcystis* genospecies have indicated a significant future change in *Microcystis* taxonomic classification^59^. However, research regarding the dynamics of microbiomes associated with different *Microcystis* morphospecies during algal blooms is practically absent.

Our use of the mNGS-IC approach and network co-occurring analysis permitted us to identify and follow four different clusters of heterotrophic bacteria associated with different *Microcystis* morphospecies. This allowed correlation of the peaks of certain *Microcystis* morphospecies at different levels of the water column with the abundances of associated microbial clusters. *Microcystis* in the natural water environment to migrate vertically, which regulates the buoyancy and changes cell density. We found that the vertically stratified distribution of cyanobacteria affected the composition and dynamics of associated microbial clusters and the heterotrophic bacteria in these clusters. Moreover, the species composition of associated microbiome clusters differed between colonial and non-colonial forms.

Accumulating evidence suggests that there may be widespread metabolic interactions between *Microcystis* and associated microbiomes^60^. The initial phase of bloom development of *Microcystis* coincided with an increase in microbial cluster numbers and included members of the ammonia-oxidizing *Nitrosomonadales* order (Cluster 1, *Methyloversatilis discipulorum*). Moreover, the decrease in the absolute abundance of *M. smithii* and *M. aeruginosa* morphospecies during the end of the first stratification period coincided with an increase in the members of the hydrocarbon-degrading gamma-proteobacteria *Xanthomonadales* (Cluster 4; *Stenotrophomonas rhizophilia*), which is similar to the observation by Gutierrez and co-authors^61^ of the structure of *Microcystis* bloom-associated microbiomes.

We hypothesized that high abundance of algicidal bacteria would coincide in time with *Microcysti*s bloom collapse, and our findings confirmed these expectations. To date, a number of algicidal bacteria have been found to be associated with cyanobacterial blooms and have been isolated and investigated^62,63^. The high diversity of anticyanobacteria reported in the literature includes more than 50 genera, with the majority being identified as members of *Pseudomonas*, *Aeromonas*, *Acinetobacter*, *Citrobacter*, and others and belonging to different Proteobacteria classes. Anticyanobacterial *Actinomycetes* include *Rhodococcus* sp., *Arthrobacter* sp., *Microbacterium,* and *Streptomyces* sp. Many *Bacteriodetes* such as *Pedobacter* sp. *Aquimarina* sp., *Firmicutes* including the *Bacillus* group, *Exiguobacterium sp.,* and *Staphylococcus* are also highly efficient in inhibiting *Microcystis* growth ^64^. In our studies, we observed a significant diversity of different *Pedobacter* sp. from the *Sphingobacteriales* order associated with the collapse of *M. wesenbergii* blooms *(Pedobacter cryoconitis, Pedobacter sp. PACM 27299, Pedobacter mongoliensis*). The seasonal dynamics of *Microcystis* morphospecies and microbial antagonists and a collapse of cyanobacterial blooms to the growth of cyanolytic bacteria have been described in a number of studies^65^, and our analysis produces highly congruent results with earlier observations that a peak of algicidal bacteria abundance coincides with or is followed by a decline in the cyanobacterial bloom^66,67^.

### Cryptophyta- and Chlorophyta-associated microbiomes

The composition of Chlorophyta- and Cryptophyta-associated microbiomes was less complex than that of the *Microcystis*-associated microbial clusters. The bloom development of Chlorophyta and Cryptophyta coincided with an increase in associated heterotrophic bacteria. The network inference analysis revealed that *Massilia*, a member of Oxalobacteriaceae, also had prevalently positive interactions with Chlorophyta, *Massilia armeniaca* thus being associated with Chlorophyta in the bottom of the vertical water column. Our results are in accordance with previous findings of positive interactions between *Massilia* and Chlorophyta^68^.

The associated microbiomes of *Cryptomonas* sp. are described by early investigators. Thus, Betaproteobacteria are abundant in many freshwater habitats and, in laboratory studies, often clustered with algae such as *Cryptomonas* sp.^69,70^. In freshwater ecosystems, one of the key bacterioplankton groups within betaproteobacteria is the genus *Limnohabitans*. Published results on co-culture have shown significant increases in the abundances of *Limnohabitans* strains in cultures of *Cryptomonas* sp. but not in cyanobacterial cultures (*Aphanizomenon* sp., *Dolichospermum* sp.)^70^. In our work, the associated *Cryptomonas* sp. microbial cluster also contained *Limnohabitans* sp. (63ED37-2). Moreover, we observed that the maximum number of the freshwater betaproteobacteria *Massilia aurea* coincided with a peak of *Cryptomonas* sp. abundance in tank D2. This confirms early observations of Salcher and co-authors^71^ reporting high numbers and growth rates of *Massilia* sp. in the presence of *Cryptomonas* sp. during co-cultivation in an artificial minimal medium.

Other members of the *Cryptomonas*-associated cluster included members of Alphaproteobacteria, from the genus *Tabrizicola (Tabrizicola pisces)* and *Gemmobacter* sp. HYN0069 (Rhodobacteriaceae) known to be associated with a high-metabolic production of phytoplankton-derived organic matter^72,73^. Another member of the identified *Cryptomonas*-associated microbiome, *Pseudobacter ginsenosidimutans* (Chitinophagaceae), has been reported to participate in plant decomposition^74^.

### Associated microbiomes and environmental parameters

*Microcystis* microbiome composition has been previously related to temperature, seasonality, *Microcysti*s morphology, and density^75–77^and changes during bloom development and degradation. The *Microcystis* microbiome bacteria have a functional potential not found in *Microcystis*^60^. We identified temperature and shifts in nutrient concentrations (phosphate) as critical factors in microbial community composition.

## CONCLUSIONS

The detailed analysis of microbiomes at the species level assessed by nanopore-based NGS in parallel with quantitative dynamics of *Microcystis* morphospecies, Chlorophyta, and Cryptophyta algal blooms revealed a complex structure of associated microbial communities. The composition of associated microbiomes changed during the seasonal bloom development and was vertically stratified for *Microcystis* morphospecies and Chlorophyta. The topological structure and dynamics of the co-occurrence network showed the complex structure and difference between colonial morphospecies and non-colonial *Microcystis*. The co-occurrence of microbial clusters with peaks of *Microcystis* collapse suggests a possible antagonistic relationship between some bacterial cluster members and *Microcystis*, which requires further research. Our investigation revealed that bloom-associated microbes deviated in response to temperature and nutrient concentrations (phosphate).

Our study highlighted that a species-level analysis is required to gain a comprehensive understanding of how microbial communities respond to algal blooms. We suggest that a combination of visualization tools (IFC) and sequencing approaches is required to evaluate the dynamics of the effect of microbiomes on planktonic behavior that is emerging to be a fundamental rule of life. This is the first analysis reporting the co-occurrence of microalgal and bacterial species based on a combination of visual (IFC) and next-generation sequencing approaches, which provides a theoretical basis for the development of detailed species-level temporal analysis of algal blooms.

## METHODS

### Collection of mesocosm samples

A schematic overview of the experimental workflow is presented in **Figure 9**. Water samples for analysis were collected at Aarhus University’s AQUACOSM Lake Mesocosm Warming Experiment (AU LMWE) facility located in Silkeborg, Denmark. The setup consisted of 24 flow-through outdoor mesocosm tanks, which combined three varying temperature regimes (ambient temperature, IPCC A2, and IPCC A2+50% climate scenarios) and two nutrient levels (high and low). Cylindrical stainless-steel mesocosm tanks are 1.9m in diameter and 1.5m in depth. In addition, the mesocosms are equipped with temperature, oxygen, and pH sensors, providing high-frequency measurements^78^. The AU LMWE 2021 experimental setup addressed the effect of temporary stratification on various aspects of ecosystem functioning by altering the mixing patterns within the mesocosm tanks. Thermal stratification was achieved by switching off mixing paddles and lifting the level of heating elements from bottom to ca. 30 – 40 cm depth. In total, 14 days of mixing were followed by 14 days of stratification during the summer period of July-August 2021. Three high nutrient tanks – D1, D2, and D3 – were the main sampling mesocosm tanks. Each tank corresponds to a temperature regime: D1 – ambient (AMB), D2 – IPCC A2, and D3 – IPCC A2+50% climate scenario. Water samples were collected for two months (once per week for eight weeks in total) with two sampling points – the surface and bottom of the tank. The bottom samples were taken first to minimize the disturbance of stratified layers within the water columns. Sequencing samples were collected in 500-mL bottles and later filtered through 0.22-um pore size GF/C filters (Whatman, USA). The filtered samples were then stored in sterile 50-mL Falcon tubes (BD Biosciences, USA) at -86°C until further analysis. Samples for IFC analysis were collected in 50-mL bottles and fixed with 1% glutaraldehyde solution. In total, 96 samples were collected and prepared.

**Figure 9.**
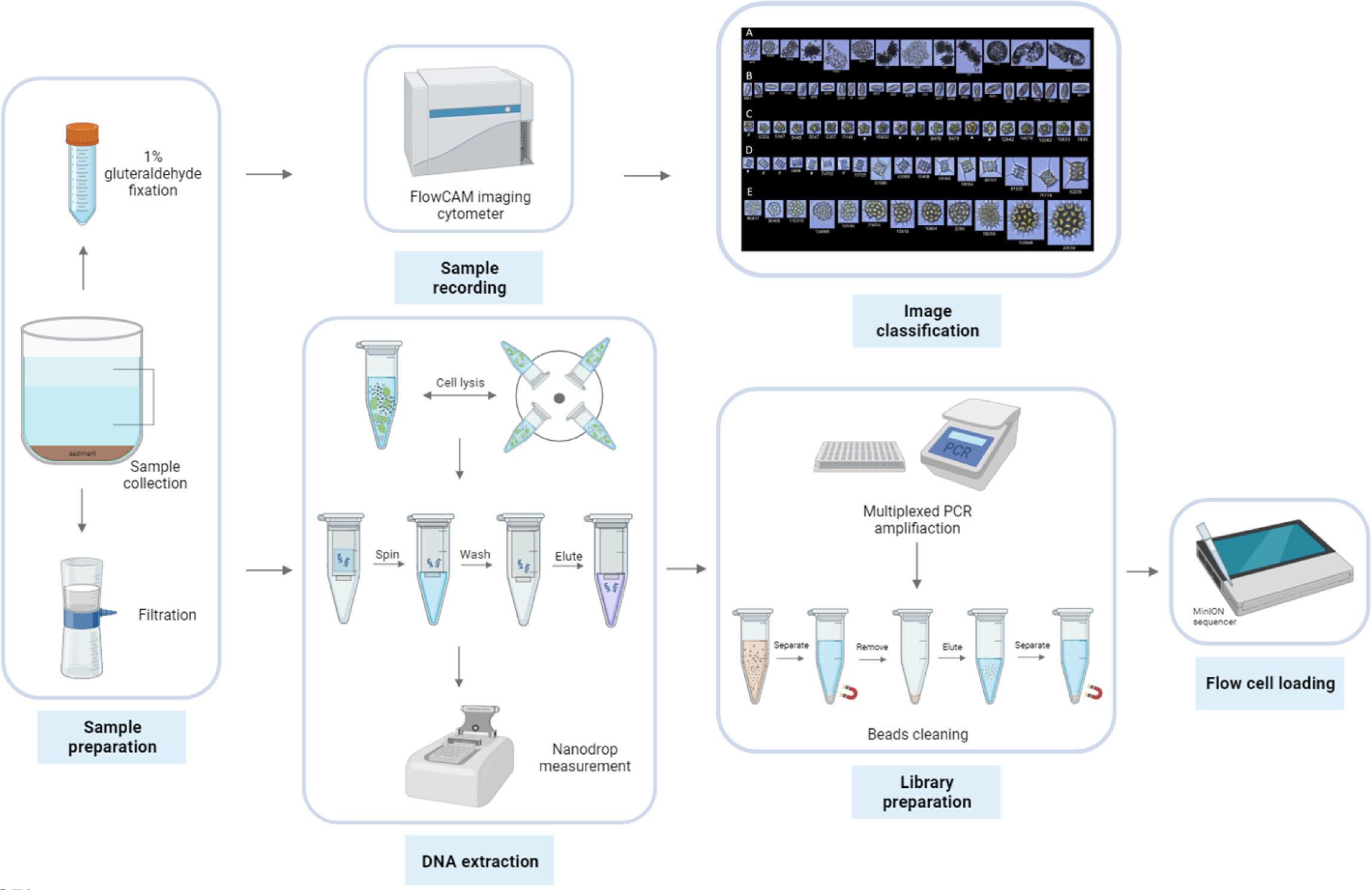
Schematic overview of experimental workflow.

### Environmental DNA extraction and sequencing library preparation

Filtration of water samples for sequencing analysis was followed by DNA extraction. For this purpose, DNEasy Power Water Kit (QIAGEN, Germany) was used to extract DNA from filters following the procedures provided by the manufacturer (with an additional lysis step with heating). In general, filtered samples were first lysed via vortexing with beads in a lysis buffer. Then, proteins and inhibitors were removed, and total DNA was captured on a spin column. After elution, 100-uL of DNA was obtained, and DNA concentrations were measured using a Nanodrop spectrophotometer (Thermo Fisher Scientific, USA). The resultant products were stored in 1.5-mL Eppendorf tubes at -20°C. DNA extraction was followed by library preparation, the first step of which involved PCR amplification. 16S Barcoding Kit 1-24 (SQK-16S024) (Oxford Nanopore Technologies, UK) was used for library preparation; it includes 24 unique barcodes and sequencing adapters necessary for multiplexed sequencing. The full-length 16S gene was amplified using universal 16S primer pair: 27F (5′-AGAGTTTGATCCTGGCTCAG-3′) and 1492R (5′-TACGGYTACCTTGTTACGACTT-3′). Library preparation was conducted according to the methodology provided by the manufacturer. The reaction mixture consisted of nuclease-free water (5 µL), input DNA (10 µL), LongAmp Hot Start Taq 2X Master Mix (25-uL), and respective 16S barcode primer (10µL). The following parameters were set for the reaction in a thermocycler: 95°C for 1 minute, followed by 25 cycles of 95°C for 20 seconds, 55°C for 30 seconds, 65°C for 2 minutes, and finally 65°C for 5 minutes. PCR products were then cleaned using AMPure XP beads (Beckman Coulter, USA), and all the barcoded samples were pooled together in a single Eppendorf tube. The final step of library preparation included the addition of sequencing adapters to the mixture of barcoded samples. Following library preparation, the MinION Mk1C device was prepared for the sequencing run. A flow cell priming kit (EXP-FLP002) was used for priming FLO-MIN106D flow cells. A newly prepared barcoded DNA library was mixed with loading beads and then loaded into the flow cell according to the manufacturer’s instructions. The sequencing run was set to about 40-46 hours.

### Bioinformatic analysis

Raw signal data were stored in FAST5 files and underwent basecalling using Guppy neural network-based basecaller integrated into the MinION Mk1C device. The obtained FASTQ files were then processed using Python commands, starting with assessing the reads and their quality using the NanoPlot package (https://github.com/wdecoster/NanoPlot). This step was followed by quality and read length filtering, with a minimum quality score set to 10, followed by removal of adapter sequences and demultiplexing of reads into respective barcodes. The demultiplexed reads were then classified using Emu taxonomic abundance estimator up to species level (https://gitlab.com/treangenlab/emu)^79,80^. The obtained data were rarefied prior to further statistical analyses.

### Imaging flow cytometry-based analysis of phytoplankton community composition

The collected samples were analyzed in parallel with IFC using a benchtop FlowCAM VS-4 imaging flow cytometer (Yokogawa Fluid Imaging, USA). Water samples fixed with 1% glutaraldehyde were analyzed with an autoimage mode using a 10X objective, followed by manual classification using VisualSpreadSheet (version 4.15.1) software. The major phytoplankton groups identified in this study are depicted in **Figure 10**.

**Figure 10.**
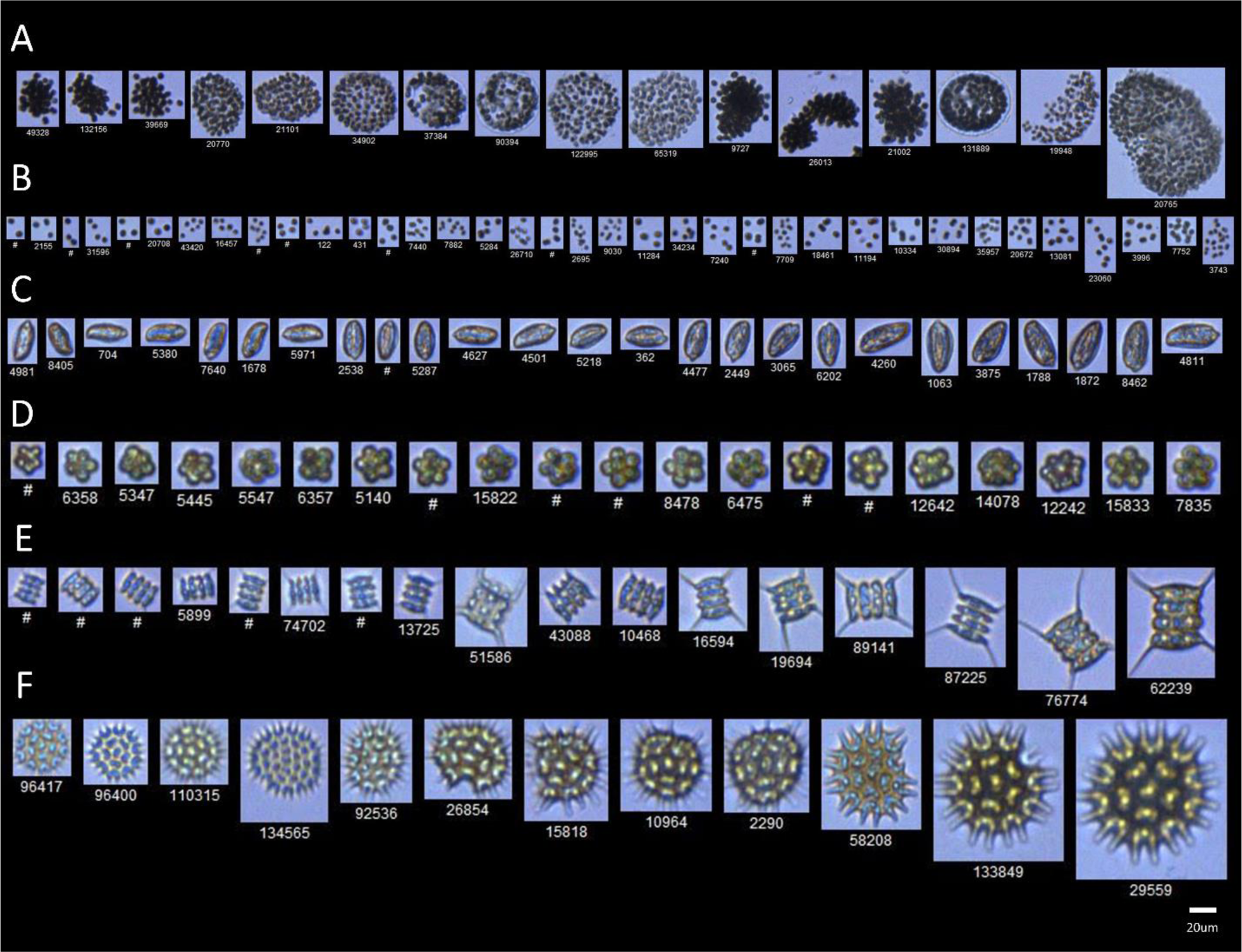
FlowCAM-based classification of major phytoplankton groups from the AU LMWE experiment. (A) Colonial *Microcystis* spp.; (B) Non-colonial small clusters (NCSC); (C) *Cryptomonas* sp.; (D) *Micractinium* sp.; (E) *Scenedesmus* sp.; (F) *Pediastrum* sp.

### Statistical analysis

The R package *vegan* (vs. 2.6-4) and Graphpad Prism software (vs. 9; Dotmatics, USA) were used for the analysis and visualization of obtained results. Diversity indices were used to characterize microbial communities within and between samples, specifically, alpha diversity metrics were applied, including observed richness and Shannon index. In addition, beta diversity metrics (Bray-Curtis dissimilarity) were assessed to quantify dissimilarity between communities and were further subjected to non-metric multidimensional scaling (NMDS). Pairwise analysis of similarities (ANOSIM) was then applied to test the significance of the differences between the obtained results. Co-occurrence networks were constructed using Pearson coefficient-based correlation matrices. Reads were first filtered to eliminate species with less than 1% relative abundance, and the obtained correlation coefficients <0.75 were then filtered out (p-value < 0.05) prior to network visualization using the R package *corrplot* (version 0.92). Networks were then visualized using the R package *igraph* (vs. 1.5.1). Canonical correlation analysis (CCA) was used to describe the relationship between microbial community composition and environmental variables (pH, temperature, oxygen level, stratification index, TP, and PO_4_-P).

## List of abbreviations

DNA: deoxyribonucleic acid
HAB: Harmful algal bloom
mNGS-IC: microbial NGS combined with imaging cytometry
NCSC: non-colonial small clusters
NGS: next-generation sequencing
IFC: imaging flow cytometry
rRNA: ribosomal ribonucleic acid
NMDS: non-metric multidimensional scaling

## Acknowledgements

This research was funded by MES Kazakhstan (grant number AP14872028) to N.S.B., by AnaEE Denmark (anaee.dk), the TÜBITAK program BIDEB2232 (project 118C250) to E.J., and by the European Commission EU H2020-INFRAIA-project (No. 731065) to T.A.D. and E.J. We are thankful to Dmitry Malashenkov from Nazarbayev University for helpful discussions and to Anne Mette Poulsen for English edition.

## Author contributions

A.M. collected and analyzed the data, contributed to the study design, wrote, and modified the manuscript, A.Z. acquired and analyzed IFC data, reviewed, and edited the manuscript, P.L. analyzed the data, reviewed, and edited the manuscript, E.J., C.S., E.E.L., and T.A.D. contributed to the funding, study design, supervision, revision, and edition of the manuscript. N.S.B. supervised the whole project, wrote, and modified the manuscript. All authors reviewed the manuscript.

## Data availability

Sequences obtained during the analysis were deposited in the National Center for Biotechnology Information (NCBI) Sequence Read Archive (SRA) under the BioProject ID PRJNA1012663. Raw data that support the findings of this study are available from corresponding author, upon reasonable request.

